# Do twitching and twitch-spindle coupling differ between N2 and N3 sleep in 6-month-olds?

**DOI:** 10.1101/2025.07.23.666445

**Authors:** Taylor G. Christiansen, Greta Sokoloff, Hailey C. Long, Olivia K. Kopp, Lydia K. Karr, Mark S. Blumberg

**Affiliations:** Department of Psychological and Brain Sciences; Iowa Neuroscience Institute University of Iowa Psychological and Brain Sciences Building 340 Iowa Ave Iowa City, Iowa 52242

**Author notes:** Corresponding author: Mark S. Blumberg, PhD Department of Psychological and Brain Sciences 340 Iowa Avenue G60 PBSP University of Iowa Iowa City, IA 52242.

**Keywords:** infancy, development, sleep spindles, myoclonic twitches, delta oscillations, sensorimotor system

## Abstract

Twitches are discrete movements that characterize REM sleep. However, recent work showed that twitches also occur during NREM sleep in human infants beginning around 3 months of age, a time when sleep spindles and the cortical delta rhythm are also emerging. Further, NREM twitches are coupled with sleep spindles, suggesting a unique contribution to sensorimotor development. Given that NREM sleep is composed of distinct substages, we investigated whether twitching and twitch-spindle coupling are differentially expressed during N2 and N3 sleep. In 6-month-old human infants (n=21; 7 females), we recorded EEG, respiration, and video during daytime sleep. We found high-intensity twitching during N2 and REM but not N3 sleep. In contrast, sleep spindles exhibited similar temporal characteristics during N2 and N3. Also, despite differences in the intensity of twitching during N2 and N3, significant twitch-spindle coupling occurred in both stages. Finally, the rate of twitching was inversely related to delta power across NREM periods. These findings suggest that although twitching occurs during REM, N2, and N3 sleep at this age, its expression is compatible with some sleep components (e.g., rapid eye movements, sleep spindles) but not others (e.g., cortical delta), highlighting the continuing need to better understand the dynamic organization of sleep and its individual components in early development.

**Statement of significance:** Twitching has long been recognized as a core component of mammalian REM sleep. However, twitching also occurs during non-REM (NREM) sleep in human infants beginning around 3 months of age. Given that NREM sleep is composed of distinct stages (including N2 and N3), we asked whether the expression of twitching and its temporal coupling with sleep spindles is different between the two stages. Even though twitching was more intense during N2 than N3, twitch-spindle coupling occurred in both stages, again suggesting a unique and likely transient functional contribution to sensorimotor development. A necessary next step is to delineate the developmental trajectory of twitching and twitch-spindle coupling beyond 6 months of age.

## Introduction

During infancy, when the brain is undergoing its most rapid development, sleep is the predominant state, with human newborns spending up to 16 hours each day asleep, evenly divided between rapid eye movement (REM) and non-REM (NREM) sleep^1^. The relatively high proportion of REM sleep in early development inspired the ontogenetic hypothesis, which posited a mechanistic relation between REM sleep—especially its phasic events—and brain development. Limb twitches are among the most prevalent phasic components of REM sleep in early development^2^. In rat pups, twitches are spontaneously generated in the brainstem^3^ to produce movements throughout the body. These movements trigger sensory responses that cascade throughout the sensorimotor system, likely refining sensorimotor circuits^2^ and contributing to the development of cerebellar-dependent internal models^4–6^.

Similar to rat pups, human infants exhibit high rates of twitching across the body during REM sleep^7^. However, in contrast to rat pups, human infants between 3 and 6 months of age also increasingly express twitches during NREM sleep^8^. The presence of twitches at high rates during NREM in infants is surprising given the apparent lack of similar reports in other mammals. Additionally, twitching exhibits distinct developmental trajectories during REM and NREM sleep: Specifically, whereas the rate of NREM twitching increases across the first six postnatal months, twitching during REM sleep occurs at a high and steady rate throughout this period.

One path to understanding the functional significance of NREM twitches is to examine the brain activity that accompanies them. For example, around the same time that NREM twitches are emerging, sleep spindles—brief thalamocortical oscillations with peak frequencies of 11-15 Hz^9^—are also emerging along the cortical sensorimotor strip^8^. NREM twitches and spindles are temporally coupled in that the probability of a spindle increases around the onset of a twitch^8^. Given the established association of spindles with learning and heightened neuroplasticity^10^, the coupling of NREM twitches with spindles suggested a unique contribution to sensorimotor development. In addition to sleep spindles, high-amplitude oscillations in the delta-frequency range (0.5-4.0 Hz; hereafter, delta) are also characteristic of NREM sleep and begin to emerge over the first few postnatal months^11^; the relation between delta and twitching has not been explicitly examined.

After sleep spindles and delta emerge, they help to distinguish the three substages of NREM sleep: Specifically, N1 is characterized by the slowing of EEG and marks the transition to sleep; N2 is characterized predominantly by the presence of sleep spindles; and N3 is characterized predominantly by the presence of cortical delta, although spindles can also be present^12^. In our previous study^8^, we did not assess whether twitching occurs differentially during NREM stages. Thus, here we characterize the rate and patterning of twitching in N2 and N3 sleep, as well as REM sleep, focusing on infants at 6 months of age when sleep spindles and delta are both clearly expressed. Our findings extend our earlier report in showing that infants twitch at higher rates during N2 and REM sleep compared with N3. Surprisingly, despite the difference in twitch rate between N2 and N3, twitch-spindle coupling is similar in both stages. Altogether, our findings highlight the complex and dynamic relations among the behavioral and electrophysiological features of sleep in early development. They also set the stage for future delineation of the full developmental trajectory of these fundamental sleep phenomena.

## Methods

### Participants

Twenty-four full-term infants, 5.35-6.99 months of age, participated in the study. Three infants were excluded from analysis because either a majority of the EEG for their session was not useable due to movement artifact, or their total sleep time was less than 5 min. Thus, the final sample comprised 21 infants (7F, 14M, 6.19 ± 0.10 months). Caregiver reports indicated that 71.4% of infants were white, 4.8% were black, and 23.8% were more than one race; additionally, 9.5% of infants were Hispanic/Latine. Of this sample, 10 were included in a previous study^8^ and 11 were newly collected. All data collection and scoring procedures are identical for all infants unless otherwise indicated. Infants and caregivers were recruited from the broader Iowa City area via email and print advertisements. All procedures were approved by the Institutional Review Board at the University of Iowa.

### Experimental design and procedure

Daytime sleep was recorded during the infant’s usual nap time. Upon arrival, infants were changed into a plain black onesie and the experimenters measured the child’s head circumference. A 128-channel HydroCel geodesic sensor net (Electrical Geodesics, Inc., Eugene, OR), sized to the child’s head, was presoaked in a body temperature potassium chloride and baby shampoo solution. After the cap was applied, electrode impedances were checked, and electrodes were adjusted until the vast majority of electrodes had impedances below 100 KΩ. Impedance values were saved at the start and the end of each sleep session. The EEG cap was connected to a Net Amps 200 Amplifier (Electrical Geodesics, Inc.). Infants slept in a supine, semi-reclined position in a baby lounger.

EEG and respiratory data were acquired at 1000 samples/s to a computer running Net Station software. A high-definition video camera (Model Q6055, Axis Communications, Lund, Sweden) provided a frontal view of the child to record movements of the eyes, face, head, and limbs. The video recording, acquired at 30 frames/s, was integrated with the EEG recording system to ensure synchrony between the data streams. When video data was exported out of Net Station software timestamps were used to synchronize physiological and video data. Recordings continued for as long as the child remained asleep. While infants slept, caregivers sat with their child in the testing room or waited in a nearby room where they could monitor the child via video.

### Behavioral coding and interobserver reliability

Sleep-wake states and twitches were visually scored from video by two raters using methods similar to those described previously^7,8^. First, two raters together reviewed videos for wake periods and the occurrence of startles and arousals. Next, periods of sleep were scored for twitches by two raters independently of one another. Using Datavyu (datavyu.org) or Net Station (Electrical Geodesics, Inc.), coders scored the video data in multiple passes to identify twitches of the face, left and right legs, and left and right arms. In addition, for the 11 participants comprising the newly collected sample, twitches were scored with more precision using Datavyu to measure twitch durations and to separately identify twitches of the toes, ankle, and legs (combining knee and hip), and twitches of the fingers, wrist, and arms (combining shoulder and elbow). Eye movements were also coded and designated as rapid eye movements (REMs) when they had durations less than 1 s. Coders did not score movements during sleep that were incidental to movements elsewhere on the body (e.g., respiration; movements of other limbs). Finally, coders were always blind to sleep stage when scoring twitches. A primary scorer was designated to score 100% of a video and a secondary scorer was designated to score at least 25% of the same video. Only periods containing both REM and NREM sleep were scored by both raters and analyzed for inter-observer reliability.

Inter-observer reliability was assessed as described previously^7,8^. Briefly, onsets and durations of twitches were determined by the two coders across all movement categories. When coders disagreed, they independently reviewed the videos around the periods of interest and, when necessary, rescored the video. After this second pass through the data, for all twitches for which each coder agreed (i.e., twitch onsets within 1 s), the onsets and durations were aligned to the primary scorer. Cohen’s kappa ranged from 0.67 to 0.99 (median: 0.87). Finally, the primary and secondary coders made a final pass together through the data record and all remaining discrepancies were resolved by mutual agreement.

### Identification of sleep stages

Two experienced raters, blind to the presence or absence of twitches, identified sleep stages in 30-s epochs based on AASM criteria^12^ for EEG activity at this age. Sleep stages were scored together by two raters and disagreements were resolved in real time to reach consensus. In addition, postural tone, REMs, and respiratory variability^13^ were used to help confirm sleep stages.

### Data analysis

#### EEG signals

Preprocessing of EEG signals was performed in MATLAB (MathWorks, Natick, MA) using EEGLab^14^. Data were band-pass filtered (0.5-70 Hz), and a 60-Hz notch filter was applied. Bad channels were visually identified and data from nearby channels were used to perform spherical interpolation. After interpolation, data were referenced to the average across all electrode sites. Data were down sampled from 1000 to 250 Hz. Periods with movement artifact were visually identified and excluded from further analysis.

Sleep spindles were detected using custom MATLAB scripts. The methods for spindle detection were described previously^8^; however, here the frequency range was expanded from 12-14 Hz to 11-15 Hz to increase sensitivity to slower frontal spindles that emerge at older ages^15^. Using a Hilbert transform of the filtered data, the amplitude of the signal was generated for artifact-free periods. The onset of a sleep spindle was marked when signal amplitude exceeded a threshold of 2x the median amplitude for at least 1 s. After this 1-s minimum criterion was met, the offset of the spindle was marked when the signal decreased below the threshold for at least 1 s. We initially reviewed all automatically detected spindles in the C3 electrode to confirm that they corresponded well with visually detected spindles in raw and filtered EEG records. In one infant we were not confident in the automatically detected spindles, so we excluded them from all spindle analyses.

Delta power was also analyzed using custom MATLAB scripts. A Hilbert transform was applied to artifact-free EEG data from the C3 electrode which was band-pass filtered in the delta frequency range (0.5-4 Hz). Delta power was calculated as the amplitude squared (µV^2^) of the transformed data at each timepoint.

#### Twitching

To obtain measures of twitch rate, the number of twitches in each body part was summed across periods of REM and NREM and divided by the total duration of each sleep stage. Twitch rates in N2 and N3 were also calculated. A total of 17 participants had REM, N2, and N3 (3 participants did not have any REM and 1 participant did not have N2). Therefore, for analysis of REM vs. NREM twitching, 18 participants were used; for the analysis of N2 vs. N3 twitching, 20 participants were used. Because twitch rates were not normally distributed (Shapiro-Wilks test, p<.05), the non-parametric Wilcoxon matched-pairs signed-ranks test was used.

Twitch duration was calculated using twitch onset and offset times as described above. For each stage of sleep, we analyzed the duration of twitches and the proportion of twitches in each body part; for the latter analysis, we divided the number of twitches in each body part by the total number of twitches for that individual. For these analyses, only newly collected data were used because these were the only data for which we had accurate twitch durations. Of the 11 participants, one did not exhibit REM sleep and was excluded from these analyses. Additionally, one participant had fewer than five twitches during N3 and was excluded from the analysis of twitching across body parts. Paired t tests were used to compare twitch durations between sleep stages, and a two-way ANOVA was used to compare the proportion of twitching across body parts and sleep stages.

Inter-twitch intervals (ITIs) were calculated as the time between successive twitches. REMs were not included. ITIs were not included if there was an intervening wake period or a transition between different stages of sleep. ITI data within each sleep stage were pooled across participants to generate log-survivor plots.

We divided twitches into those occurring as part of a burst or as single events. Bursts were defined as a series of at least two twitches with <0.5 s between successive twitch onsets. A twitch burst ended when the onset of the next twitch occurred at an interval >0.5 s. Therefore, single twitch events occurred >0.5 s after the preceding twitch and >0.5 s before the subsequent twitch. Burst rate was calculated by dividing the total number of bursts during each sleep stage by the total time spent in each sleep stage. Burst rates had non-normal distributions (Shapiro-Wilks test, p<.05); therefore, the non-parametric Friedman test was used, and a Conover test was used for pairwise comparisons using PMCMRplus package in R^16^. We also analyzed the proportion of twitches occurring in bursts in each sleep stage, calculated by dividing the number of twitches occurring in bursts by the total number of twitches. Only participants with at least 10 twitches in each sleep stage were used in these analyses; 12 participants met this criterion.

#### Sleep spindles

Spindle rate was calculated by dividing the total number of spindles detected at each electrode site by the total number of minutes spent in artifact-free N2 or N3. Topoplots of spindle rates excluded a subset of 46 channels around the neck and ears as these channels frequently contained movement artifact. Because the distributions of spindle frequencies in the F3, C3, and P3 electrodes were nearly identical, subsequent analyses focused on spindles at the C3 electrode. Spindle amplitude was considered as the maximum voltage (µV) of the Hilbert-transformed signal for each spindle waveform; sleep spindle frequency (Hz) was also determined. Mean spindle amplitude and mean spindle frequency during N2 and N3 were computed for each participant. In addition to the one participant excluded from all spindle analyses, three participants were excluded because they had fewer than 2 min of artifact-free N2 or N3. Thus, the final spindle analyses included 17 participants. For comparing spindle rate, amplitude, and frequency, paired t tests were used.

Inter-spindle intervals (ISIs) were calculated as the time between successive sleep spindles at the C3 electrode. ISIs were excluded if there was an intervening wake period or a transition between stages of sleep. ISI data within each sleep stage were pooled across participants to generate log-survivor plots.

#### Association between delta power and twitching

We analyzed the relations between delta power and twitches during NREM sleep. To conduct these analyses, mean delta power and number of twitches were calculated for each 1-min window of artifact-free NREM sleep. First, for each infant, we conducted a linear regression relating delta power to the number of twitches. Next, we used a multilevel linear mixed-effects analysis to evaluate whether delta power predicted variations in twitches across infants. Models were estimated using the lmer function within the lme4 package in R^17^. Intraclass correlations (ICCs) indicated that 57% of the variance in delta power was due to between-infant differences. Therefore, delta power was separated into two predictors: (1) a between-infant measure representing an infant’s mean delta power across all NREM windows (MeanDelta_i_ for each infant *i*), and (2) a within-infant measure representing an infant’s delta power deviation in that window from their individual mean (WithinDelta_ti_ for each timepoint *t* for each infant *i*). This approach allows us to examine whether twitching was related to differences in average delta power across infants (between-infant effect) versus fluctuations across time in an individual infant’s mean delta power (within-infant effect)^18^. All delta measures were divided by 100 such that a one unit increase in MeanDelta_i_ or WithinDelta_ti_ indicates a 100 µV^2^ increase in delta. The MeanDelta_i_ is centered around the grand mean of delta such that when MeanDelta_i_ is zero, the infant mean delta power is 1100 µV^2^. The WithinDelta_ti_ was centered around the individual mean: a higher or lower value represents a higher or lower amount of delta in that window relative to the infant’s mean.

The equation for the model is as follows:

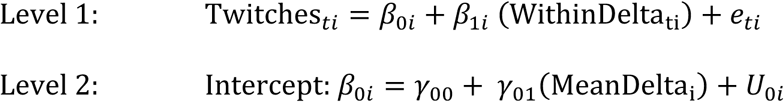

Within − Person Delta: 𝛽_1*i*_ = 𝛾_10_

In the Level 1 model, the number of twitches across time (Twitches_ti_) is a function of an individual intercept (β_0i_), the within-person effect of delta power (β_1i_), and the residual number of twitches (e_ti_) at timepoint *t* for infant *i*. In the Level 2 model, the individual intercept (β_0i_) is a function of a fixed intercept (γ_00_), the main effect of between-infant delta power (γ_01_), and an individual-specific random intercept (U_0i_). The individual effects of within-infant delta power (β_1i_) are equivalent to the fixed effect (γ_10_). Multilevel models were estimated using restricted maximum likelihood (REML) estimation^19,20^.

#### Coupling between twitches and sleep spindles

For each participant, we identified the total number of spindles that co-occurred with a twitch. For each spindle, if at least one twitch occurred within the onset and offset of the spindle, it was classified as co-occurring with a twitch. We calculated the probability of a spindle given a twitch (i.e., P(Spindle | Twitch)), by dividing the number of twitches co-occurring with a spindle by the total number of twitches. P(Spindle | Twitch) was calculated during NREM sleep at each electrode site across the scalp. Then, P(Spindle | Twitch) during NREM sleep was averaged across 19 central electrodes (including C3, C4, and Cz) and 14 frontal electrodes (including F3, F4, and Fz). We next computed perievent histograms to assess the probability of a sleep spindle occurring in the 40-s period around the onset of a twitch (i.e., ± 20 s). Perievent histograms were computed at each electrode and then averaged across central and frontal sites. Shuffled data were used to compute 95% confidence intervals (p < 0.05) at each time point.

The next set of analyses were performed to compare N2 and N3 at the C3 electrode only. First, we calculated P(Spindle | Twitch) as described above. Second, we calculated the inverse, that is, the probability of a twitch given a spindle (i.e., P(Twitch | Spindle)) by dividing the number of spindles co-occurring with twitches by the total number of spindles. For these analyses, nine participants had very brief N2 or N3 durations; thus, we only included participants in this analysis if they had at least 10 spindles and 10 twitches in both N2 and N3, which left 12 participants for analysis.

Perievent histograms for the probability of a spindle in relation to twitch onset were computed as described above. Perievent histograms were also computed to assess the probability of a twitch occurring in the 40-s period around the midpoint of a C3 spindle. Only those participants with at least 10 spindles and 10 twitches in either N2 or N3 were included in this analysis, leaving 14 participants in N2 and 16 participants in N3.

Finally, to examine if the P(Twitch | Spindle) and P(Spindle | Twitch) occurred at above-chance levels, onset times for the trigger events were shuffled using a Monte Carlo procedure. For instance, for P(Twitch | Spindle), the onset times of the spindles (i.e., the triggers) were shuffled 1000 times in a manner that retained the original distribution of inter-spindle intervals; only the order of the intervals was randomized (a parallel procedure was used when the triggers were twitches). For each of the 1000 permutations of randomized data, the method described above for calculating mean probabilities was repeated and a grand mean for each plot was calculated. The probabilities from the shuffled data were considered the expected probabilities. To test for differences in the observed and expected probabilities during NREM sleep between central and frontal electrodes, a two-way repeated-measures ANOVA was performed. Similarly, to test for differences in the observed and expected probabilities at the C3 electrode during N2 and N3, a two-way repeated-measures ANOVA was performed.

#### Statistical analyses

All statistical analyses were conducted in SPSS Statistics 29.0 (IBM, Armonk, NY) unless otherwise specified. Before conducting t tests or ANOVAs, data were tested for normality using the Shapiro-Wilks tests. When normality assumptions were violated, non-parametric tests were used. Sphericity was tested using Mauchly’s test. When sphericity was violated, a Huynh-Feldt correction was applied to the degrees of freedom. Multiple pairwise comparisons were adjusted using the Dunn-Šidák correction. The central tendency of the data is reported as the mean, and dispersion as SEM. The measure of effect size for ANOVAs is represented as partial eta-squared.

## Results

### The rate of twitching does not differ between NREM and REM sleep

We recorded electrophysiological and behavioral data during daytime sleep in 6-month-old infants (n=21). Approximately half (n=10) of the data were published previously^8^ and the remainder were newly collected for this study. Across all sessions, infants slept an average of 26.73 ± 2.76 min, comprising 5.44 ± 1.15 min of REM sleep, 4.87 ± 0.64 min of N2 sleep, and 16.25 ± 2.40 min of N3 sleep.

As expected from our previous report, twitches occurred during NREM sleep, that is, in the presence of sleep spindles and delta oscillations when REMs were absent (**Figure 1**). We also observed twitches in the presence of REMs, indicative of REM sleep. The rate of twitching during NREM (9.13 ± 1.16 twitches/min) was not significantly different than the rate during REM (13.70 ± 2.68 twitches/min; *Z* = -1.24; **Figure 2A_1_**). The duration of individual twitches also did not differ between NREM (0.44 ± 0.04 s) and REM (0.51 ± 0.05 s; *t*(9) = -1.74; **Figure 2A_2_**). As described previously ^8^, a higher proportion of twitches occurred in some body parts (e.g., fingers and toes) than other body parts (e.g., arms and legs; **Figure 2A_3_**); ANOVA revealed a significant main effect of body part (*F*(2.45, 22.04) = 12.72, *p* < .001, ηD^)^ = 0.59), but neither the main effect of sleep stage (*F*(1, 9) = .51) nor the body part x sleep stage interaction was significant (*F*(2.61, 23.46) = 1.95), indicating that twitch patterning across the body was similar in REM and NREM. Thus, at 6 months of age, there are no discernible differences in the characteristics of twitches during NREM and REM sleep.

**Figure 1.**
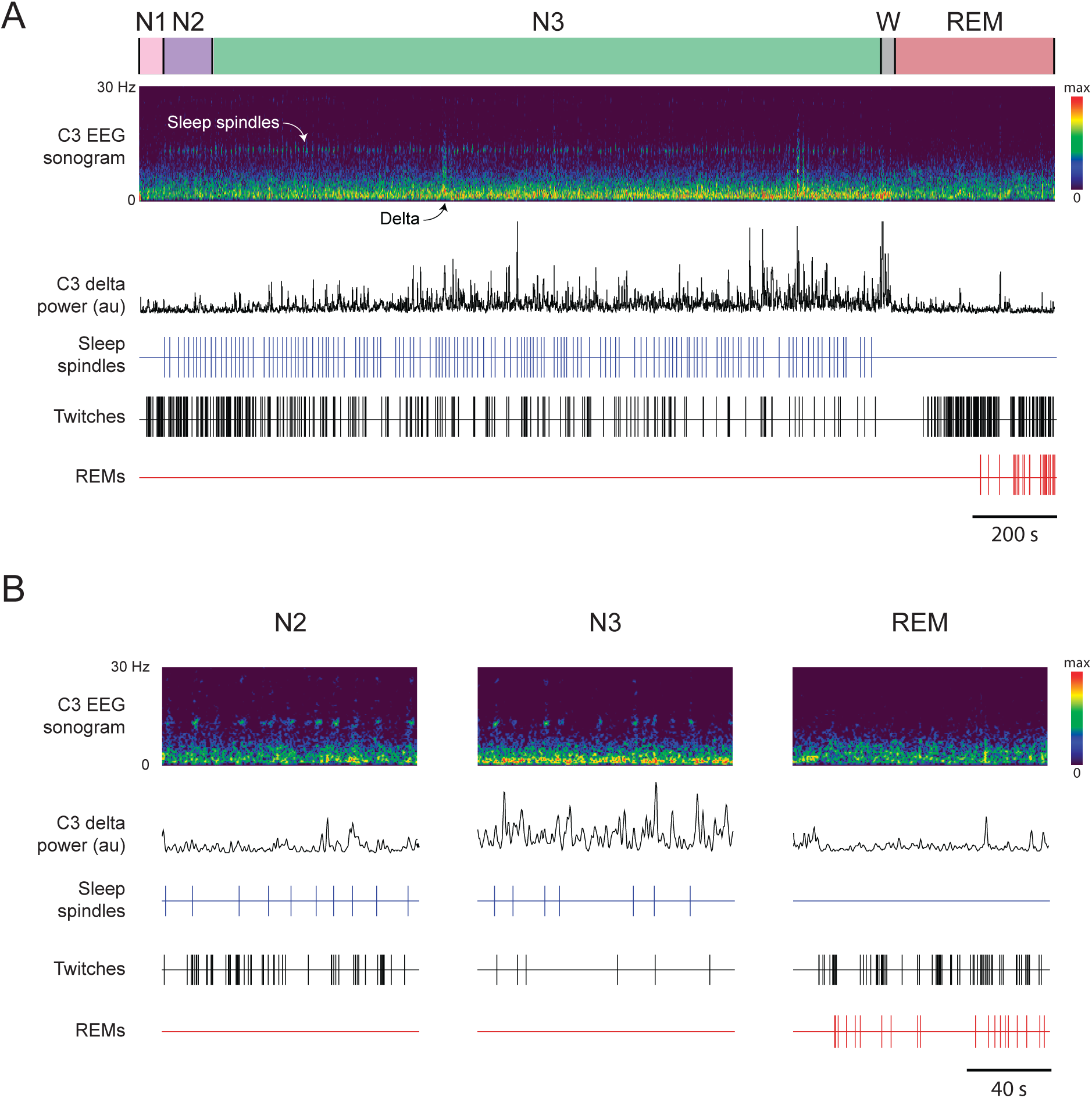
Representative behavioral and EEG data in a 6-month-old human infant. **(A)** Continuous 37.5-min record of N1 (pink), N2 (purple), N3 (green), wake (W, grey) and REM (red). EEG data are from the C3 electrode site and are represented as a sonogram (0-30 Hz) and delta power (0.5-4 Hz, black trace). The onset of spindles (blue ticks), limb twitches (black ticks), and REMs (red ticks) are also shown. **(B)** Same infant and data presentation as in (A) but for 2-minute periods of N2, N3, and REM sleep.

**Figure 2.**
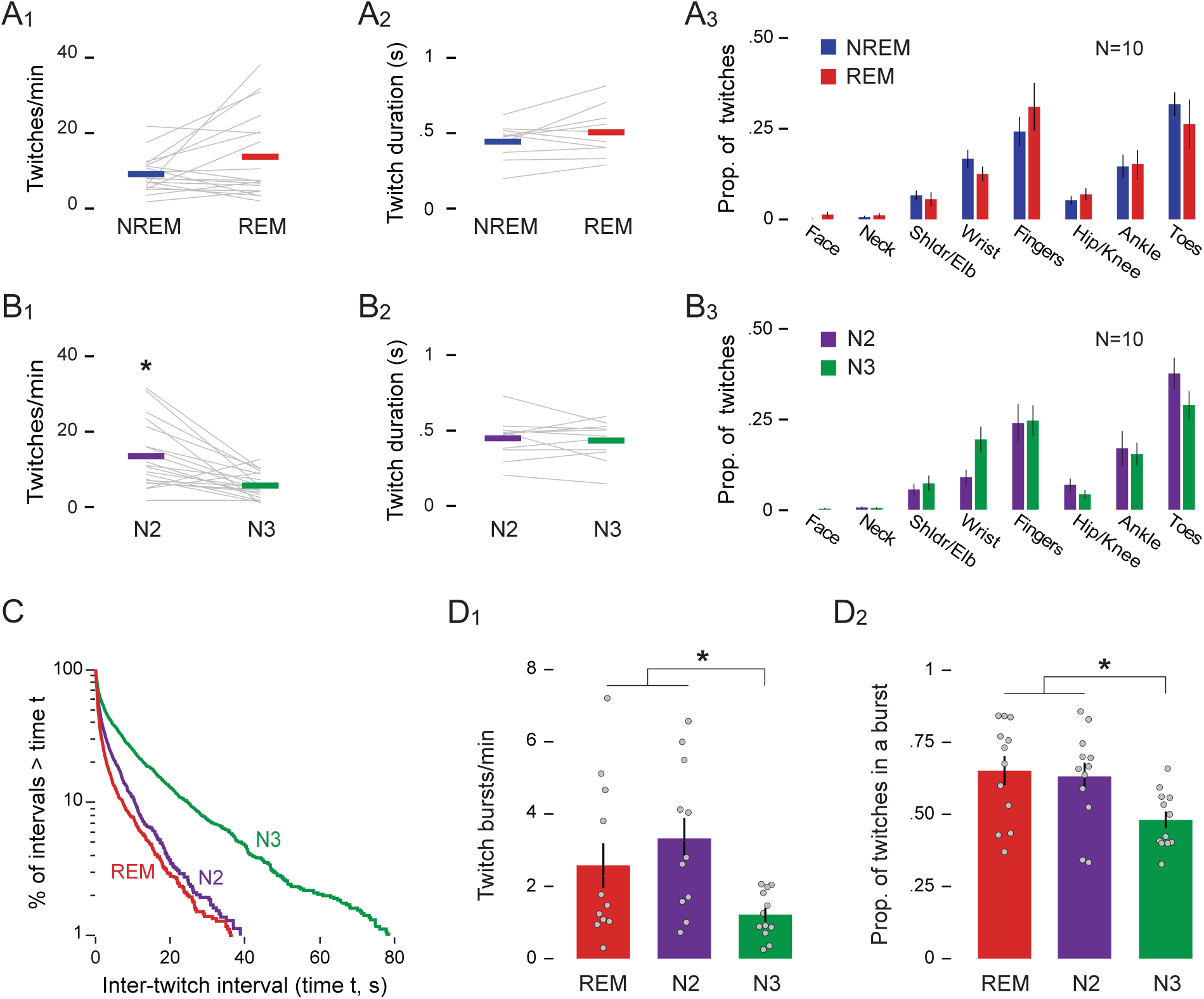
Characteristics of twitching during REM and NREM sleep. (A_1_) Mean rate of twitching (twitches/min), **(A_2_)** mean duration of twitches (s), and **(A_3_**) mean (±SEM) proportion of twitching across various body parts during NREM (blue) and REM (red) sleep. Grey lines show data for individual infants. **(B_1-3_)** Same as in (A_1-3_) but for N2 (green) and N3 (purple) sleep. **(C)** Log-survivor plot of inter-twitch intervals during periods of REM, N2, and N3 sleep. ITIs were pooled across sleep sessions: REM = 1795 twitches; N2 = 1239 twitches; N3 = 2060 twitches. **(D_1_)** Mean (±SEM) twitch-burst rate (bursts/min) and **(D_2_)** mean (±SEM) proportion of twitches occurring in a burst during REM, N2, and N3 sleep. Grey dots show data for individual infants. * p < .05

### Twitch characteristics differ between N2 and N3 sleep

Although the rate of twitching did not differ between NREM and REM sleep, it did differ significantly within NREM between N2 (13.45 ± 1.95 twitches/min) and N3 (5.68 ± 0.77 twitches/min; Z = -3.21, p = 0.001; **Figure 2A_1_**). However, the duration of individual twitches did not differ significantly between N2 (0.45 ± 0.04 s) and N3 (0.43 ± 0.04 s; t(10) = 0.50; **Figure 2B_2_**). Also, although the distribution of twitches across the body was not uniform, it was similar in N2 and N3; specifically, there was a significant main effect of body part (*F*(2.57, 23.12) = 16.20, *p* < .001, ηD^)^ = 0.64), but neither the main effect of sleep stage (*F*(1, 9) = 1.00) nor the body part x sleep stage interaction was significant (*F*(3.59, 32.34) = 2.75; **Figure 2B_3_**).

To further illustrate differences in the rate of twitching across sleep stages, we created log-survivor plots of inter-twitch intervals, pooled across infants, during REM, N2, and N3 (**Figure 2C**). For log-survivor plots, the slopes of the distributions are proportional to the rate of twitching, with steeper slopes indicating higher rates. Similar to previous reports of the temporal organization of twitching in human infants^7^ and infant rats^21^, the slopes of the distributions in all three states were steepest at the shortest intervals, indicating that many twitches occurred in rapid succession (i.e., in bursts). Also, it is evident from the slopes of the log-survivor plots that the rates of twitching were very similar during REM and N2, and both were higher than N3.

The log-survivor plots supported our visual assessment of the raw data (see **Figure 1B**) that twitching in all three stages, particularly in REM and N2, often occurs in bursts. To further analyze this phenomenon, we defined a burst as a series of two or more twitches in which each inter-twitch interval was less than 0.5 s. The rate of bursts was significantly different among the three sleep stages (𝜒² (2) = 12.17, p = .002; **Figure 2D_1_**), with bursts occurring at more than twice the rate during REM (3.33 ± 0.56 bursts/min) and N2 (2.58 ± 0.62 bursts/min) than during N3 (1.22 ± 0.19 bursts/min). Additionally, a higher proportion of twitches was part of a burst during REM (0.65 ± 0.05) and N2 (0.63 ± 0.05) than during N3 (0.48 ± 0.03; *F*(2, 22) = 8.768, p= .002, ηD^)^ = .443; **Figure 2D_2_**). In summary, the differences in the rate and burstiness of twitching lead us to conclude that twitching was more intense during N2 and REM than during N3.

### Sleep-spindle characteristics do not differ between N2 and N3

To account for the differences in twitching between N2 and N3, we first considered the possibility that they are related to a lower rate of sleep spindles in N3, as is characteristic in adults^9,22,23^. However, we found similar rates of sleep spindles across the scalp between N2 and N3, with sleep spindles occurring primarily in the frontal and central electrodes (**Figure 3A**). Also, the frequency of sleep spindles, with peaks at 13-13.5 Hz, did not differ across the scalp (**Figure 3B**). Given these findings, we focused the remaining analyses on spindles at one central electrode cite, C3.

**Figure 3.**
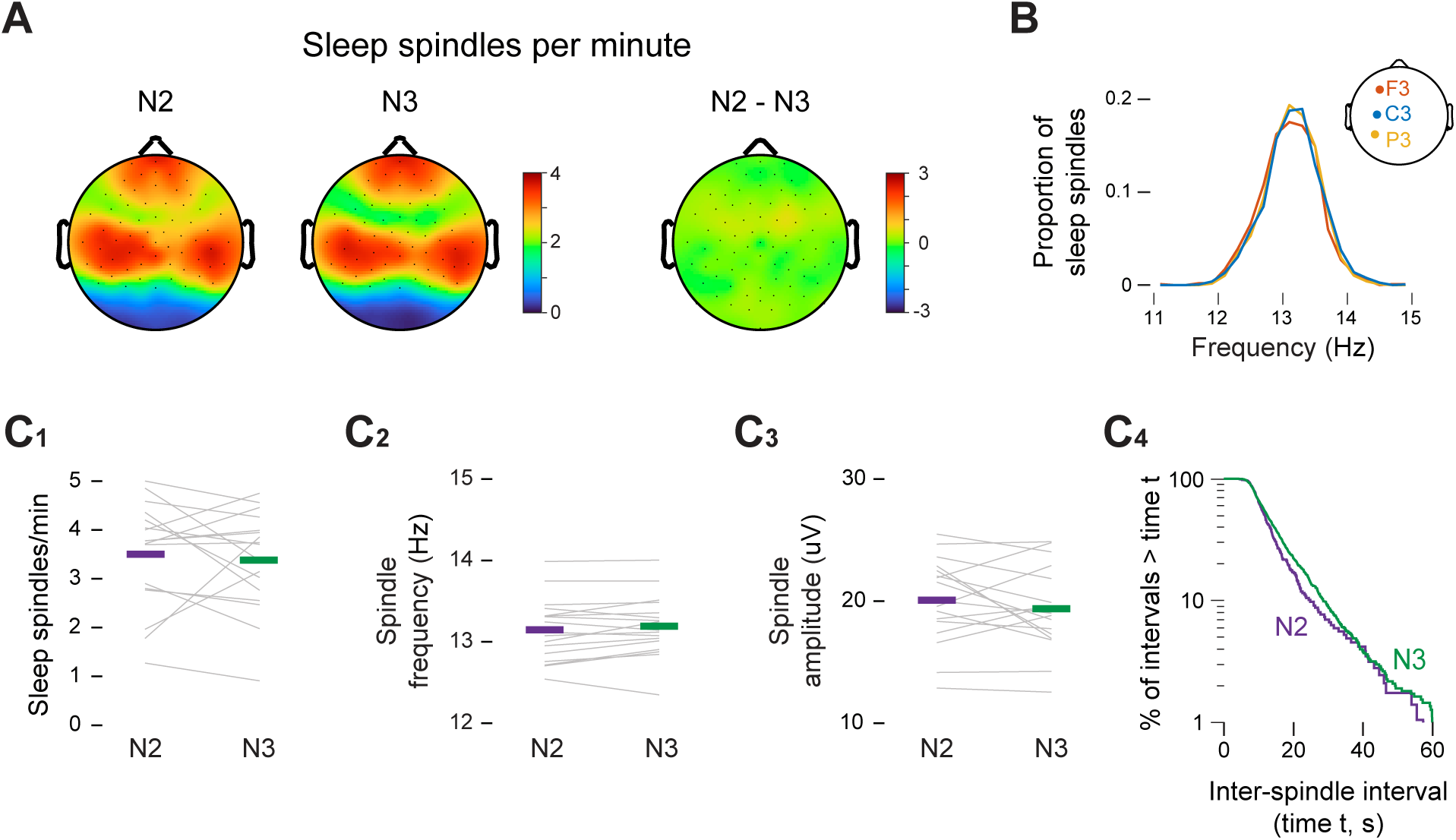
Characteristics of spindles do not differ between N2 and N3 sleep. **(A)** Topoplots showing the mean spindle rate (spindles/min) across the scalp during N2 and N3 sleep (left) and the difference between N2 and N3 (right). **(B)** Frequency distribution of sleep-spindle frequencies pooled across participants at the F3 (orange), C3 (blue), and P3 (yellow) electrodes. **(C_1_)** Mean spindle rate (spindles/min), **(C_2_)** mean spindle frequency (Hz), and **(C_3_)** mean spindle amplitude (µV) at the C3 electrode during N2 (green) and N3 (purple) sleep. Grey lines show data for individual infants. **(C_4_)** Log-survivor plot of inter-spindle intervals (ISIs) in the C3 electrode during N2 (green) and N3 (purple) sleep. ISIs were pooled across sleep sessions: N2 = 286 spindles; N3 = 1104 spindles.

In the C3 electrode there was no significant difference in the rate of spindles between N2 (3.50 ± 0.26 spindles/min) and N3 (3.38 ± 0.25 spindles/min; t(16) = 0.52; **Figure 3C_1_**). Also, there were no significant differences in peak spindle frequency (N2: 13.15 ± 0.09 Hz; N3: 13.19 ± 0.09 Hz; t(16)= -1.53; **Figure 3C_2_**), or spindle amplitude (N2: 20.08 ± 0.85 µV; N3: 19.38 ± 0.84 µV; t(16) = 1.10, **Figure 3C_3_**). Finally, log-survivor plots of pooled inter-spindle intervals show nearly overlapping distributions for N2 and N3 (**Figure 3 C_4_**). These findings argue against the possibility that spindles are related to the lower rate of twitching in N3.

### Inverse relations between twitching and delta power

We next considered whether the presence of cortical delta during N3 is associated with its lower rate of twitching. For each infant, we calculated linear regressions relating mean delta power to the number of twitches across all 1-min windows of artifact-free NREM sleep. Within individual infants, we found inverse correlations between delta power and twitching (mean R^2^ = 0.29 ± 0.07; **Figure 4A**).

**Figure 4.**
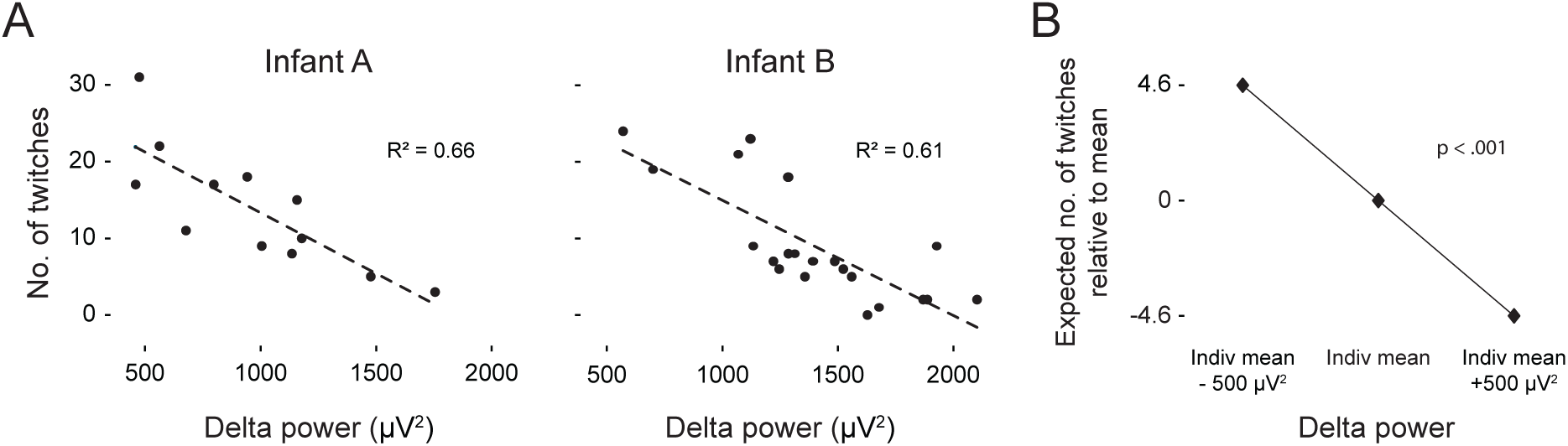
Twitches are inversely related to delta power. **(A)** Representative linear regressions from two infants showing the number of twitches in relation to mean delta power (µV^2^) across 1-min windows of artifact-free NREM sleep. **(B)** Multilevel linear mixed-effects analysis of the effect of within-infant changes in delta power on the expected number of twitches within each 1-min window.

Next, a multilevel linear mixed-effect model was used to evaluate whether delta power predicted variation in the rate of twitching across infants (**Figure 4B**). There was a significant within-infant effect of delta, such that for every 500 µV² increase/decrease in delta power during a 1-min window, the number of twitches in that window was expected to be 4.6 twitches higher/lower than that infant’s average (t(288.61) = -10.04, p < 0.001; **Table 1**). In contrast, between-infant variation in delta power did not significantly predict the rate of twitching: Infants with higher average delta power did not exhibit lower rates of twitching. In summary, these findings suggest that within periods of NREM, higher delta power, as occurs during N3, is incompatible with higher rates of twitching.

**Table 1.** Results of mixed-effect model of the association of within-and between-infant fluctuations in delta power with the rate of twitching during NREM sleep.

| Effect | Estimate | Std Err | DF | t Value | p Value |
| --- | --- | --- | --- | --- | --- |
| Intercept | 6.07 | 0.83 | 15.43 | 7.29 | <.001 |
| Within-Infant Delta | -0.93 | 0.09 | 288.61 | -10.04 | <.001 |
| Between-Infant Delta | -0.01 | 0.25 | 16.28 | -0.04 | .971 |

### Both frontal and central sleep spindles are temporally coupled with twitches

We previously reported that the probability of sleep spindles in central electrodes increases significantly around the time of a NREM twitch^8^. However, sleep spindles occur at high rates in both frontal and central electrodes, so here, we compared the temporal relationship between both frontal and central spindles in relation to NREM twitches. We first assessed the probability of a sleep spindle across all electrodes given the occurrence of a twitch, which revealed spindle probabilities are highest in the central and frontal electrodes (**Figure 5A**). Overall, the probabilities were similar in a group of central (0.16± 0.02) and frontal electrodes (0.18 ± 0.02; **Figure 5B**). The observed probability of a spindle given a twitch was significantly greater than expected by chance across both electrode areas (F(1, 19) = 59.30, p < 0.001, ηD^)^= 0.80). In contrast, neither the main effect of electrode area (F(1, 19) = 1.09) nor the electrode area x observed-expected interaction (F(1, 19) = 1.51) was significant. Additionally, in both central and frontal electrodes the probability of a spindle exceeded the 95% confidence intervals in the period surrounding a twitch (**Figure 5C**). Thus, given that a twitch occurred, sleep spindles in central and frontal electrodes are equally likely to co-occur with that twitch.

**Figure 5.**
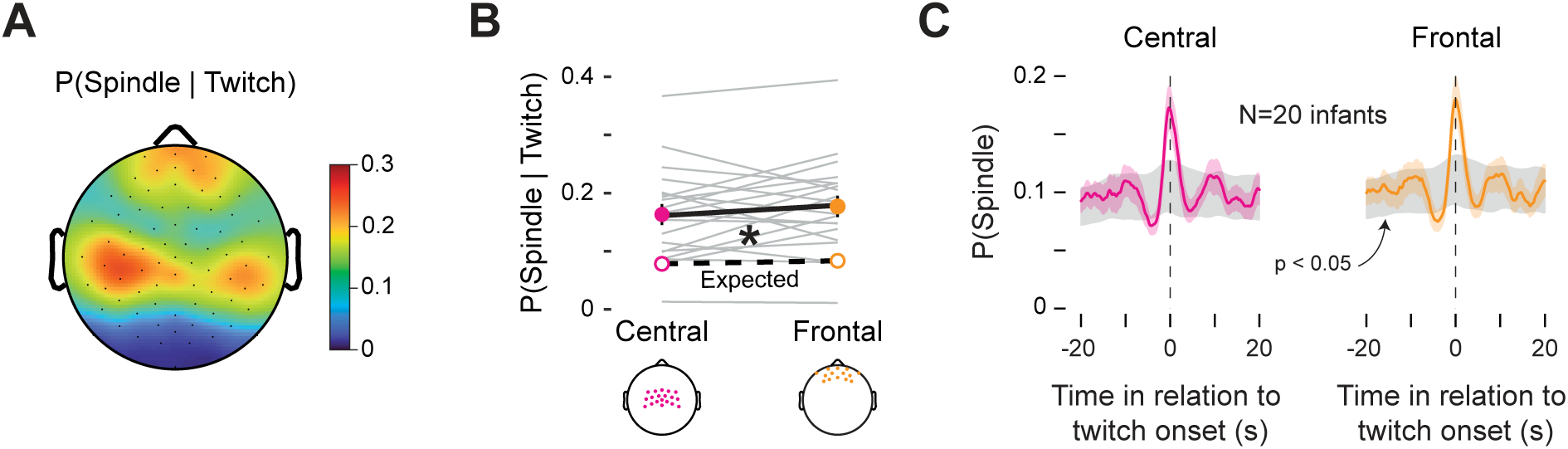
Spindles in frontal and central electrodes are coupled with twitches during NREM sleep. **(A)** Topoplot of the mean probability of a sleep spindle given the occurrence of a twitch (i.e. P(Spindle | Twitch)). **(B)** Mean (±SEM) P(Spindle | Twitch) calculated across central electrodes (pink solid circle) and frontal electrodes (orange solid circle). The central and frontal electrode sites included in these analyses are visualized below. Grey lines show observed data for individual infants. Mean expected probabilities (unfilled circles) based on shuffled data are also shown. (**C)** Perievent histograms of mean P(Spindle | Twitch) for central (left; pink) and frontal (right; orange) electrodes. Each plot shows mean (±SEM) probabilities. The grey shaded regions denote 95% confidence intervals based on shuffled data.

### Sleep spindles are temporally coupled with twitches in both N2 and N3

Next, we analyzed the temporal coupling of spindle and twitches separately during N2 and N3. Given that spindle twitch coupling was not different between central and frontal electrodes, we focused the remaining analyses on spindles occurring in the C3 electrode site. We first assessed the probability of a sleep spindle given the occurrence of a twitch (**Figure 6A_1_**). During N2 and N3, the probability of a spindle exceeded the 95% confidence intervals in the period surrounding a twitch (**Figure 6A_2_**). Overall, the probabilities were similar during N2 (0.29 ± 0.02) and N3 (0.26 ± 0.03; **Figure 6A_3_**). The observed probability of a spindle given a twitch was significantly greater than expected by chance across both sleep stages (F(1, 11) = 77.72, p < 0.001, ηD^)^= 0.88). In contrast, neither the main effect of sleep stage (F(1, 11) = 1.93) nor the sleep stage x observed-expected interaction (F(1, 11) = 0.11) was significant. Thus, given that a twitch occurred, a sleep spindle was equally likely to co-occur during N2 and N3.

**Figure 6.**
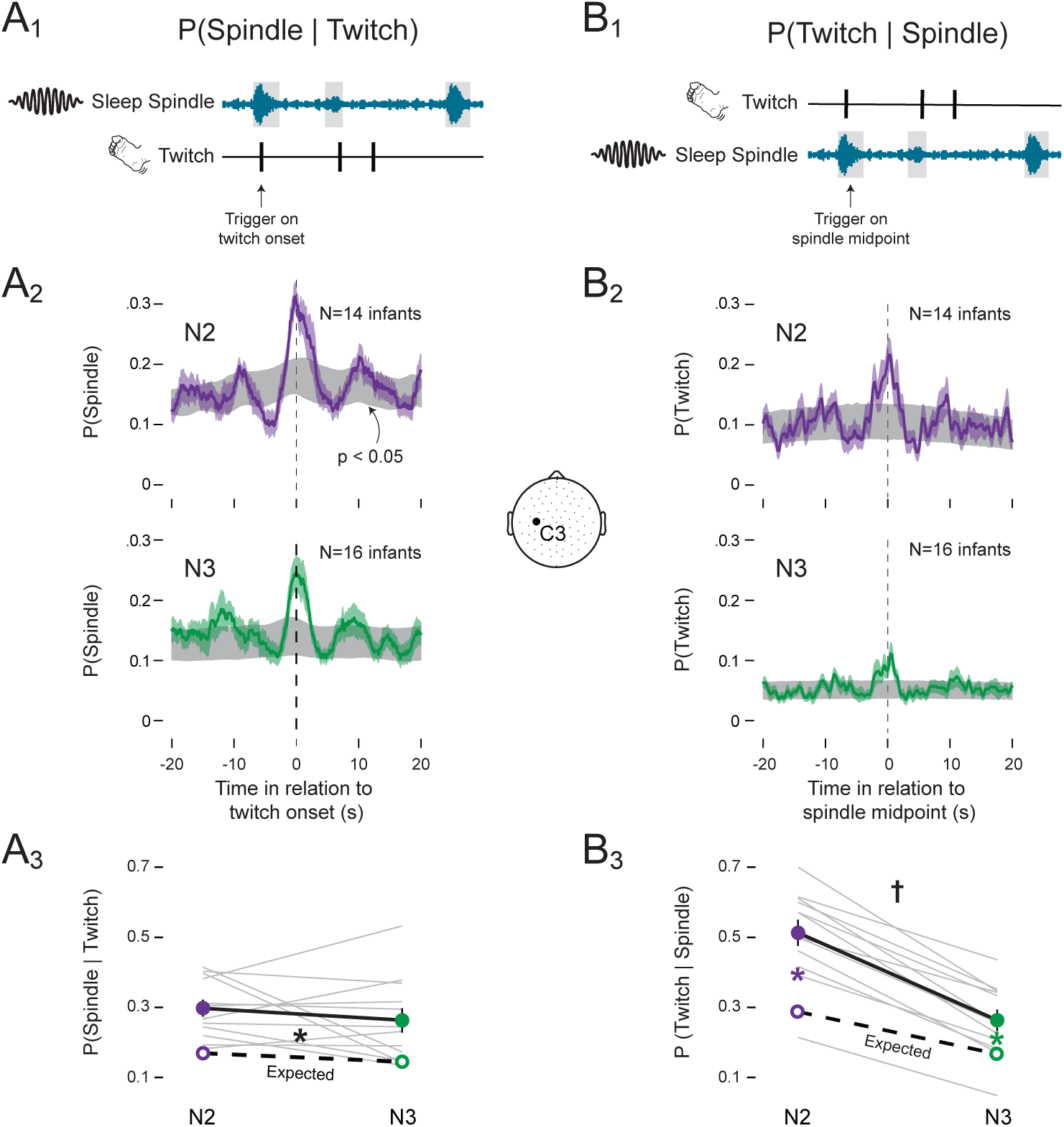
Spindles and twitches are coupled in N2 and N3 sleep. (A_1_) Illustration of our method for assessing the probability of a sleep spindle given a twitch: P(Spindle | Twitch). For every twitch, we determined whether a spindle (from onset to offset) overlapped with that twitch. **(A_2_)** Perievent histograms showing the P(Spindle | Twitch) during N2 (top; purple) and N3 (bottom; green) sleep. Each plot shows mean probabilities with SEM (shaded values). For each plot, 95% confidence intervals based on shuffled data are shown in grey. **(A_3_)** Mean (±SEM) P(Spindle | Twitch) during N2 (purple solid circle) and N3 (green solid circle). Grey lines show observed data for individual infants. Expected mean probabilities (unfilled circles) based on shuffled data are also shown (expected probabilities for individual infants are not shown). **(B_1-3_)** Same as in (A_1-3_) but for the probability of a twitch given a sleep spindle: P(Twitch | Spindle). Black * significant main effect (p<.05) between observed and expected values. Colored * significant observed and expected probability differences in N2(purple) and N3(green) (p<0.05). † Significant interaction (p<.05) between sleep state and observed-expected probability differences.

Next, we inverted the analysis and measured the probability of a twitch given the occurrence of a sleep spindle (**Figure 6B_1_**). During N2 and N3, the probability of a twitch exceeded the 95% confidence intervals in the period surrounding a sleep spindle, although the peak during N3 was lower (**Figure 6B_2_)**. Overall, the probability was 0.51 ± 0.04 during N2, compared to only 0.26 ± 0.03 during N3 (**Figure 6B_3_)**, consistent with the significantly lower rate of twitching in N3 (see **Figure 2B_1_**). Given a sleep spindle, ANOVA revealed that a twitch was significantly more likely to co-occur during N2 than N3, as evidenced by the significant sleep stage x observed-expected interaction (*F*(1, 11)= 13.483, p=.004, ηD^)^ = 0.55, **Figure 6B_3_**). Post hoc tests revealed that for both N2 and N3, the observed probabilities were significantly greater than expected by chance (N2: t(11) = 6.609, p < 0.001; N3: t(11) = 4.640, p < 0.001). Thus, although the lower rate of twitching during N3 reduced opportunities for spindles to co-occur with twitches, twitch-spindle coupling was still greater than expected by chance.

## Discussion

Building on the surprising discovery that twitches emerge postnatally during NREM sleep and are temporally coupled with sleep spindles^8^, here we delved deeper to determine whether these phenomena are expressed differentially during N2 and N3. The first main finding is that twitching during N2 and REM is more intense than during N3, reflecting its significantly higher rate and tendency to occur in bursts. Second, the reduced rate of twitching during N3 was inversely related to cortical delta power, suggesting an incompatibility between delta and twitching. Finally, despite differences in twitching between N2 and N3, twitch-spindle coupling occurred in both stages. Overall, these findings highlight the changing organization of sleep over development and suggest that twitch-spindle coupling, which is not expected to persist into adulthood, makes a transient functional contribution to early sensorimotor development.

The main limitations of this study are the relatively small and geographically limited sample size and the focus on a single age. We chose 6 months because it is the oldest age investigated in our prior study^8^, and the infants at that age exhibit distinct N2 and N3. Thus, it was a good choice for extending our previous study to examine the issues of differential twitching and twitch-spindle coupling during N2 and N3. Moreover, in addition to reanalyzing the data from the infant participants in that previous study, we also collected new data to double the size of the prior dataset. Next, we are building on this detailed assessment at 6 months of age to delineate the developmental trajectories of twitching and twitch-spindle coupling over subsequent months and years.

### Why is the rate of twitching lower in N3?

Twitches occurred at half the rate during N3 compared to N2 (and REM). Why? Based on known differences in spindle rates between N2 and N3 in adults^9,22,23^, and the twitch-spindle coupling during NREM in infants^8^, we considered the possibility that, at least during NREM, differences in spindle rate are associated with differences in twitching.

However, we found that spindle rates in N2 and N3 did not differ. This finding is consistent with an earlier study in infants at 4 and 6 months of age^24^. Indeed, one recent study in 6-8-month-olds reported a higher spindle rate during N3^25^. Therefore, spindles cannot explain the differential rate of twitching in N2 and N3.

Given that the cortical delta rhythm distinguishes N3 from both N2 and REM, perhaps the presence of delta is incompatible with twitching (**Figure 7**). Consistent with this idea, we found that within-infant fluctuations in delta power were negatively associated with NREM twitch rate. Assuming this association has a causal basis, through what mechanism might cortical delta influence twitching? One possibility is that cortical delta directly interferes with the production of twitches by motor cortex; however, based on the current understanding of twitch production during REM sleep in infant^3,26^ and adult^27,28^ animals, twitches are produced in the brainstem, not the cortex. Thus, if cortical delta were able to somehow modulate the production of twitching, it would have to do so by somehow modulating subcortical networks.

**Figure 7.**
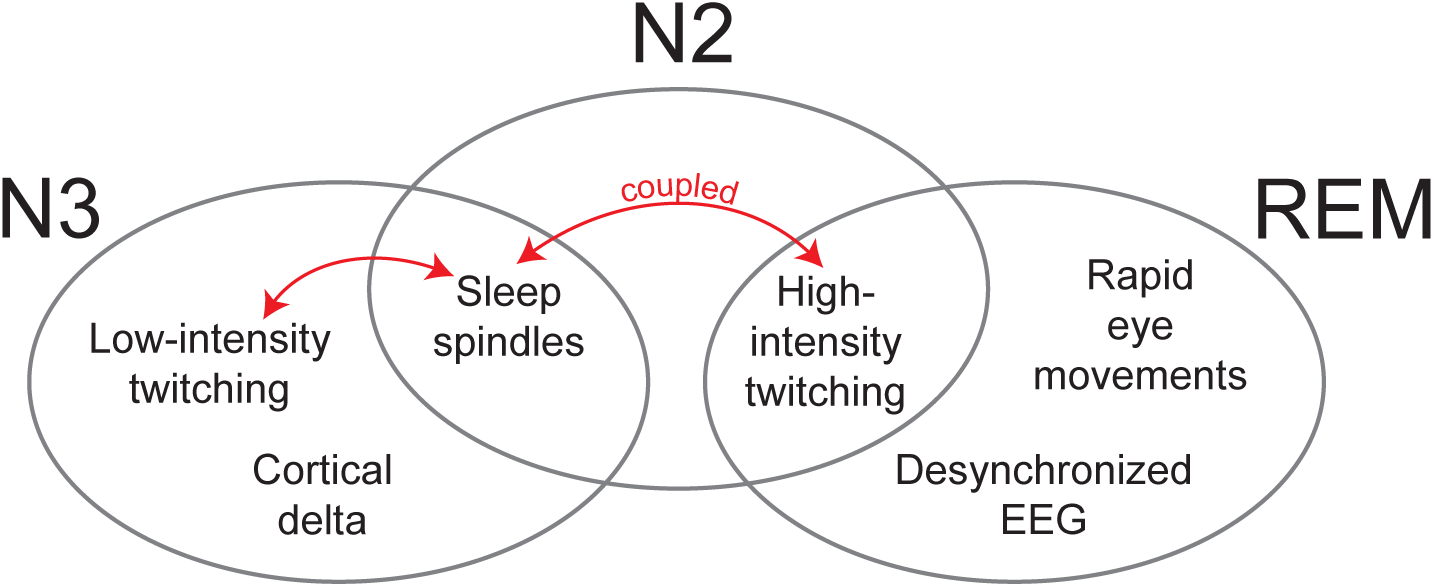
Organization of sleep components of REM, N2, and N3 at 6 months of age. Venn diagram to illustrate the sleep-stage distribution of twitching (high and low intensity), sleep spindles, rapid eye movements, and cortical EEG activity (i.e., delta, desynchronized). The red double arrows indicate the coupling of twitches and spindles that occurs in N2 and N3.

An alternative explanation is that cortical delta is a proxy for other associated processes in the brainstem that are causally related to twitching. For example, it was recently reported in 12-day-old rats, the age when cortical delta is first emerging, that delta oscillations also occur within a medullary structure, the parafacial zone^29^, that is implicated in the regulation of NREM sleep^30^. Not only was spiking activity within the parafacial zone phase-locked to the locally produced delta, but the delta rhythms in the cortex and parafacial zone were coherent. Thus, it may be that the presence of cortical delta signifies a brain state that is incompatible with the production of twitching. Testing this idea in human infants may not currently be feasible, but future assessments of the developmental trajectory of twitching in relation to cortical activity during N2 and N3 may help to inform our understanding of possible contributing mechanisms.

### Twitch-spindle coupling occurs similarly in frontal and central spindles

We investigated how spindle characteristics differed between central and frontal electrodes. In adults, sleep spindles recorded at frontal electrodes occur on average at slower frequencies (10-12 Hz) than at central electrodes (13-15 Hz)^9,31^. These frequency differences appear to reflect distinct mechanisms and functions, with central spindles being more tightly linked to memory consolidation^32–35^. Importantly, this distinction between central and frontal spindles is absent in early infancy and only emerges gradually between 6 months and 2 years of age^36,37^. Consistent with earlier findings^25,36^, in our participants across all electrode locations, including central and frontal electrodes, spindles exhibited peak frequencies of 13-13.5 Hz. Also, sleep spindles in central and frontal electrodes were similarly coupled with twitches. Beyond 6 months of age, as fast and slow spindles begin to differentiate, a thorough understanding of the trajectory of twitch-spindle coupling in sensorimotor and frontal cortex may provide valuable insight into the functional development of these structures.

### Twitch-spindle coupling occurs during N2 and N3

Twitch-spindle coupling is a robust phenomenon; for example, at least half of the sleep spindles during N2 co-occurred with a twitch. Thus, given the established association between sleep spindles and neuroplasticity^9,10,38^, twitch-spindle coupling may make a unique functional contribution to typical sensorimotor development. Moreover, twitch-spindle coupling may open new avenues for understanding atypical development, including neurodevelopmental disorders that are marked by deficits in sensorimotor function and sleep spindles. Sleep-spindle deficits have been well documented in individuals with schizophrenia^39–41^, autism^42–44^, cerebral palsy^45^, and other neurodevelopmental disorders^46^. Sensorimotor problems are also common features of many of the same neurodevelopmental disorders^47–52^. Therefore, twitch-spindle coupling provides new opportunities for exploring how sleep spindles contribute to neurodevelopmental disorders.

### Why have NREM twitches in infants not been reported previously?

There are a variety of methodological and conceptual explanations for why previous investigators appear to have overlooked the presence of NREM twitches in infants. First, although twitching has long been identified as a core component of REM sleep, there have been few systematic investigations of the spatiotemporal properties of twitching in humans at any age. Instead, twitching has typically been assessed in a binary fashion— as either being present or absent^53–57^. Moreover, in our experience, the careful assessment of twitches from video is still the most accurate way to detect twitching across the body; accelerometry and other similar methods fail to detect the vast majority of twitches, and time-lapse video misses twitching altogether. Thus, one can easily understand why investigators who are not interested in twitching per se might fail to notice its occurrence outside of REM sleep, especially given the strong association of twitching with REM sleep. Without systematic behavioral assessments of twitching in relation to EEG and other electrophysiological signals, periods of twitching with NREM-related components would either be ignored, misclassified as REM sleep, or designated as indeterminate sleep.

The considerations above are only magnified by the likelihood that twitching during NREM sleep, at least at rates similar to those during REM sleep, may only exist within a narrow developmental window. In adults, there are reports of twitching—particularly finger twitching—during NREM sleep, but it occurs at much lower rates than during REM sleep^58–60^, and twitch-spindle coupling is only a recently discovered phenomenon^8^. Thus, the present findings—made possible by real-time, coordinated behavioral and electrophysiological assessments of sleep—lay the foundation for a full explication of NREM twitching and twitch-spindle coupling over the infant and toddler years.

### Conclusions

By detailing the spatiotemporal structure of twitching in relation to other sleep components at 6 months of age, this study reveals new aspects of the dynamic organization of infant sleep. The dynamic concept of sleep organization was first introduced to account for the dramatic changes in the sleep of premature infants^61^. But it is clear that equally dramatic changes continue throughout the postnatal period, including the emergence of NREM twitching and twitch-spindle coupling around 3 months of age^8^. We extended our earlier findings here by showing that twitching is differentially expressed during N2 and N3 at 6 months of age. Also, although twitching occurs across all stages of sleep at this age, some sleep components (e.g., REMs, sleep spindles) are compatible with high-intensity twitching, while other components (e.g., delta) are not (**Figure 6**). Looked at another way, N2 shares characteristics with both REM (i.e., high-intensity twitching) and N3 (i.e., sleep spindles). Such findings defy conventional adult-centric assumptions of sleep organization and underscore the difficulty of cleanly differentiating sleep stages in infancy when what we describe as “sleep” is itself changing over time.

Research in adults tells us where sleep organization is headed, but not how it gets there. Notably, we do not expect the high rates of NREM twitching or twitch-spindle coupling to persist into adulthood. Thus, it is likely that both of these aspects of infant sleep are transient, not unlike many other transient developmental features^62^. Thus, looking beyond 6 months of age, we expect that a full description of the developmental trajectory of these seemingly transient NREM phenomena will provide critical clues regarding their neural substrates and functional significance for the developing infant.

## Acknowledgments.

This research was supported by grants from the National Institute of Child Health and Human Development (R01-HD104616; R21-HD095153) to M.S.B. We thank Jimmy Dooley for help with MATLAB scripts, and Becky Yu, Hayley Chappel, and Daylan Carney for help with behavioral scoring.

## Disclosure Statement

Financial disclosure: none. Non-financial disclosure: none.

## Data Availability

By the time of publication, coded data and a subset of videos will be shared on Databrary. Custom MATLAB scripts will be shared on Github.

## Notes

### Competing Interest Statement

The authors have declared no competing interest.

### Summary of Updates

This manuscript was revised and resubmitted after peer review.

